# Pharmacophore-driven Identification of N-Methyl-D-Receptor Antagonists as Potent Neuroprotective Agents Validated Using *In-Vivo* Studies

**DOI:** 10.1101/415646

**Authors:** Mukta Sharma, Anupama Mittal, Aarti Singh, Ashwin K. Jainarayanan, Swapnil Sharma, Sarvesh Paliwal

**Author notes:** **Corresponding Author’s mailing address and email ID:** Mukta Sharma, Department of Pharmacy, Banasthali University, P.O. Banasthali, Rajasthan-304022, Email I.D.

## Abstract

Alzheimer’s disease (AD), the most widespread cause of dementia is delineated by progressive cognitive impairment in the elderly people. During its progression, N-Methyl-D-Aspartate receptor antagonists are known to play a key role in the mechanisms of learning and memory. Extensive side effects alongside other effects on learning and memory have limited the therapeutic significance of various blockers and antagonists of the NMDA receptor. In this study, we identify potential compounds targeted against NMDA. In order to reveal the essential structural features for NMDA receptor, three-dimensional pharmacophore models are constructed based on a set of known NMDA inhibitors. This is followed by virtual screening which results in novel chemical compounds having the potential to inhibit NMDA. The lead compounds are then subjected to molecular docking and assessed by a scoring function, which results in two compounds with high Libdock scores. These compounds also show interactions with important residues at the active site. The compounds are shortlisted on the basis of high estimated activity, fit values, LibDock score, no violation to Lipinski’s and availability for procuring.

Of the shortlisted compounds, one compound satisfying the entire aforementioned criterion is further tested using *in-vivo* studies on mice with the help of an eight-arm radial maze. The pharmacophore-based virtual screening protocol presented in this study pave the way forward to address the unmet medical need of Alzheimer disease.

## Introduction

Alzheimer’s disease (AD) is a gradual neurodegenerative disease accompanied by mental deterioration with large amount of neuronal loss. AD is mainly characterized by the presence of neurofibrillary tangles, amyloid plaques and the degeneration of neurons in weak brain areas such as the hippocampus and the neocortex. Although fundamental mechanisms of AD neuropathology are not well understood, upcoming evidences imply that altered NMDA receptor function and enhanced NMDA receptor-mediated excitotoxicity may add to both functional and pathological irregularity of AD. The N-Methyl-D-Aspartate receptor antagonists play a crucial role in the mechanisms of learning and memory which are the most basic cognitive functions to be influenced during the normal aging process^1^. Generally the NMDA receptor is a heteromeric ligand-gated ion channel in the Central Nervous System and are present pre-synaptically albeit at a high density postsynaptically^2,3^. Recently, it has also been discovered that NMDA receptors exhibit tetrameric assemblies. Each subunit comprises of a huge extracellular amino terminal domains^4^. In general, there are several sites on the NMDA receptor that can vigorously inhibit, activate or enhance the functioning of the receptor. These receptors are attached to high conductance cationic channels that are permeable to K^+^, Na^+^ and Ca^2+^. The property of high Ca^2+^ permeability make NMDA receptors apt for their function in intervening synaptic plasticity, which bring about development and learning processes^5^. However, extensive side effects and effects on learning and memory have limited the therapeutic significance of various blockers and antagonists of the NMDA receptor. Literature evidence suggests that N N’-Diarylguanidine derivatives^6^ which are the specific antagonists of the NMDA receptor may have the ability to show a better potential as therapeutic reagents due to their high affinity and improved selectivity.

Therefore, in the present work, we identify novel and structurally diverse NMDA receptor antagonists through a well defined sequential *in-silico* virtual screening protocol followed by the biological evaluation of the lead compound.

In this paper, we report a lead compound, HTS 00987, which is thoroughly validated by computational tools and tested by *in-vivo* studies for the treatment of AD. This compound can further be subjected to clinical trials for its development as a novel drug to treat AD. In order to achieve that, we use a 4-phase approach (Ligand-Based Drug Designing, Virtual Screening, Molecular Docking and Biological Evaluation) to discover novel leads as neuroprotective agents. In the first phase, we use the ligand-based drug designing methods which utilize three-dimensional properties of the ligands to predict the biological activities. The pharmacophore model with the highest correlation and the best RMS fit are then subjected to pharmacophore mapping and various validation studies. In the second phase, we apply the validated models for the database search in order to retrieve most potent compound. The lead compounds with good fit values, estimated activity, drug-likeness and docking score are checked for novelty by employing pairwise tanimoto similarity indices using “Find Similar Molecules by Fingerprint” protocol in Discovery Studio. In this study, all the lead compounds show low Tanimoto similarity indices to all the structures of known NMDA receptors antagonists validating their uniqueness^7^. The third phase entails molecular docking studies succeeded by evaluating the retrieved potent lead compounds for neuroprotective activity.

## Materials and methods

### Pharmacophore Modeling

Pharmacophore modeling is one of the most powerful and efficient approaches to identify a unique scaffold which can be generated based on ligands. Pharmacophore model was generated with an endeavor to represent the collection of key features which are vital for biological activity^8^. The HYPOGEN method was applied for the ligand based pharmacophore modeling which utilizes the biological activity values of the compounds in the training set to build the hypothesis by using Discovery Studio V2.0 software (Accelrys Inc., San Diego, CA, USA).

The “BEST” algorithm was employed to construct 255 conformers per molecule with an energy threshold of 20 kcal/mol. Excluded volumes were also considered during pharmacophore generation^9^.

### Test and training set preparation

For hypothesis generation, training set molecules ought to satisfy a definite set of laws as it should be broadly occupied by structurally varied compounds (minimum 16) along with the most active compounds which have to be necessarily included in the training set and their activity range must lie between three to five orders of magnitude. For the present study, a set of 40 different compounds with biological activity values (IC_50_) ranging between 8 nM and 13,000 nM were chosen as a “training set” to produce the hypotheses. In order to substantiate the hypothesis, the test set was organized in a similar manner to training set. The “Test set” included 19 structurally diverse compounds as of the training set by a broad range of biological activity values. Chem Draw 8.0 was exploited to outline the two-dimensional (2D) chemical structures of all compounds which were changed later into 3D structures using DS. A maximum numbers of “255” conformations were produced for every compound by employing the “BEST” conformation method for model generation which is based on CHARMm force field to guarantee the energy-minimized conformation of every molecule with an energy threshold of 20 kcal/mol. All the possible conformations of the compounds were considered and were used for hypothesis generation using DS^10,11^.

### Pharmacophore generation using HYPOGEN

A 3D QSAR pharmacophore model was produced by evaluating the biological activity values of shortlisted compounds in the training set. The feature mapping module from DS was employed to choose the significant chemical features for hypothesis generation. Whilst generating hypotheses, HBD and HY features were chosen dependig on the training set compounds with a minimum of 0 to a maximum of 5 features^12,13^. The ‘Uncertainty’ values for the 59 compounds in the training set were 3, and values for additional parameters were kept constant.

Afterward, pharmacophore models were computed using 3D QSAR Pharmacophore module and high scoring hypotheses were collected^14^. The top scoring hypotheses were selected on the basis of correl, rms, weight, configuration, fixed cost, null cost, and total cost values^15^, ^16^.

### Assessment of pharmacophore quality and database screening

To assess the quality of the pharmacophore, three different approaches were used. Primarily, cross-validation was carried out by rearranging the data with Fischer’s randomization test. The results validated that the hypotheses generated from the training set are sound^16^. Also, the calculation of test set suggested a correlation value of 0.65 between the experimental and predicted activities. Finally, an external test set was used along with a well known marketed drug namely memantine for the validation.

## Methods for validating pharmacophores

### 1. Fisher’s Cross-Validation Test

To assess the statistical significance of the generated models, we used Fisher’s randomization test with a motivation to examine the activity data associated with the training set. The training sets (randomized) are used to generate hypotheses using the similar pharmacophoric features and parameters. If the data sets (randomized) yields pharmacophoric models with a better cost values, rms and correlation, then the original hypothesis is believed to have been generated by chance. With the help of the Cat Scramble program which is available in the Catalyst HypoGen module, the biological activities of the molecules in the training set were analysed and the resultant training sets were employed for the HypoGen runs. Therefore, all parameters were used in a similar fashionto the initial HypoGen calculation. It was observed that, at a 99% confidence level, Cat Scramble created 99 spreadsheets^17,18^.

### 2. Internal test set Validation

An internal test set comprised of 19 compounds, displaying diverse activity classes and different functional groups was used to determine the predictive ability of the derived model. The predictive value of test set was estimated in terms of the squared correlation coefficient (r^2^). The prediction of desired activity was calculated on the basis of feature mapping of test set molecules onto the developed pharmacophore and their respective fit values.

The best mapped compound was compared with least mapped compound. The fit value depends on the number of pharmacophore features that efficiently superimpose to the appropriate chemical moieties, the weight of the relevant hypothesis features spheres, the distance between the center of a particular pharmacophoric sphere and the center of the associated superimposed chemical moiety of the fitted compound^19,20^.

### 3. External test set Validation

The generated pharmacophore was also validated by employing a structurally different external set of NMDA inhibitors. The actual activity of these compounds ranged from 8 to 3000 nM. The selected ten external test set compounds were mapped onto the developed pharmacophore model. It was also validated by mapping a well known marketed drug “memantine” which is an NMDA based inhibitor as an additional quality check method. The mapping pattern (fit value) and difference between estimated and actual activity for all the compounds were thoroughly examined^21^.

## Virtual screening

Virtual screening, a broadly used tool for identification of leads in silico has helped pharmaceutical industry to increase the chances of designing medicines in a lesser time^22^. Though a thoroughly validated pharmacophore model contains the significant chemical features accountable for biological activities of potential drugs, it can be used as a 3D query to run a database search. As mentioned before, the best hypothesis (Hypo1) was utilized as a 3D query for obtaining effective molecules from the chemical databases such as NCI and Maybridge. A total of 250 conformers with a maximum energy tolerance of 20 kcal/mol were generated for every molecule in the database with the help of “best conformer generation method”. To begin with, the compounds were sorted by Lipinski’s “Rule of five” that sets the criteria for drug-like properties^23^. Drug-likeness is a measure to distinguish novel lead compounds by screening various available structural libraries. According to this rule, “a compound is not suitable for further studies if “MW > 500, log P > 5, hydrogen bond donors > 5, and hydrogen bond acceptors > 10”. Furthermore, the molecules which showed a complete feature mapping over the query pharmacophore model was selected as a hit^24^.

To accomplish this process, two database searching options (Fast/Flexible and Best/Flexible search) are used. Of these two, the “Best/Flexible search” resulted in improved results throughout database screening, therefore, we completed all database searching experiments using the “Best/Flexible search” approach. The selected pharmacophore model was employed to screen the databases for the compounds that satisfied all five features of the selected pharmacophore. The fit values were also calculated for selected hits on the basis of the pharmacophoric features and their deviation from the centers of the features. Compounds with higher fit values indicate good matches. The compounds that qualified all of these screening tests were shortlisted for molecular docking studies.

## Molecular Docking Studies

The lead compounds obtained from ligand based database search were subjected to docking studies to check the type of molecular interactions^25^. Molecular Docking studies were carried out using the programme LibDocker which is a molecular dynamics simulated annealing based algorithm and available as an extension of DS V.2.0^26^. The crystal structure of NMDARs obtained from PDB was used for docking. The receptor protein was verified by assessing its valency, missing hydrogens and any structural disorders like connectivity and bond orders. The selected protein was divided into the protein and ligand part. The protein part was labeled as a receptor molecule while the ligand was utilized to delineate the binding site of approximately 9 Angstroms on the receptor molecule. The chemical structure of the test compound was sketched and subjected to energy minimization in order to get the highly stable structure for molecular docking studies ^26, 27, 28^. Based on present coordinates, marketed drug and test compounds were employed to molecular docking studies and all the parameters were set to their default values. At the end, all possible interaction modes were analyzed on the basis of Libdock Scores.

## Biological Evaluation

Since, there is no known cure for Alzheimer’s disease till date, it has become one of the biggest challenges in the field of medicine. On the basis of high estimated activities, fit values, LibDock score, violation to Lipinski’s and availability for procuring, one compound (HTS 00987) was finally subjected to *in-vivo* studies on mice using an eight-arm radial maze^29,30^.

This study was carried out in two different sets in order to assess the effect of extended treatment of diazepam-induced amnesia^31,32^. In the first set, the treatment was provided for 7 days to mice whereas in the second set the treatment was provided for 14 days. All experiments performed were approved by the Institutional animal ethical committee. Young albino mice weighing 25-30g, provided by the CCSHAU - Hisar were used for behavioral testing. This study was performed according to the OECD 432 guidelines. The mice were divided into two sets of two groups each. The duration of treatment for animals of set I and set II was 7 and 14 days respectively. Mice were delivered two trials through the day. Every trial started with the careful cleaning of the maze by using 70% ethanol. Two arms (no. 1 and no. 3) of the arm maize were baited with food. Mice were carefully placed at the center of the octagon and were allowed an open choice which was followed by recording the selection of baited arm. The arms of the maze were not re-baited, so the first entry into the baited arm was noted as the correct choice^33,34^. After three weeks of training, the mice were given CMC (carboxymethylcellulose, 0.5%), memantine (5mg/Kg), and HTS 00987 (5mg/Kg) which were administered twice (10 am and 7 pm) at the same time on each day orally for a period of 7 and 14 days. The experimental study was designed such that the impact of CMC, memantine and HTS 00987 could be assessed after 7 and 14 days against diazepam-induced amnesia. At the end of treatment, all the animals were subjected to diazepam (1mg/kg i.p.) 60 minutes later the drug administration, excluding the first group of every set which toiled as a vehicle control. The cognitive parameters were assessed 30 minutes later the administration of the inducing agent (Diazepam) using an eight-arm radial maze, which corresponds to learning^35^. The lead compound HTS 00987 was evaluated for behavioral studies using diazepam-induced amnesia in mice model. On the 8th and 15th day respectively, mice were placed in the center of the octagon. In order to investigate the effect of control/CMC, memantine and HTS 00987 on wistar albino mice, five parameters were examined: number of entries in the baited arm, duration in the baited arm, %correct choice, RME (Reference memory error) and WME (Working memory error) using ***ALL MAZE* software**^36^. Percent choice was estimated by halving the number of correct entries by the total number of entries in the baited arm and multiplied by 100. An entrance into an unbaited arm was counted as reference memory error and re-entry in the baited arm was regarded as working memory error^37^. Average values for each variable were calculated for week 1 (1-7 days) and week 2 (8-14 days) and one way ANOVA was performed on the data for each week.

## Results and Discussion

### HYPOGEN model for N N’-Diarylguanidine inhibitors

In the current study, we collected a set of 40 diverse compounds with their corresponding biological activities from the literature. The training set with extensive diversity includes the N N’-Diarylguanidine inhibitors of N-Methyl-D-Aspartate Receptor Antagonists. Biological activities and structures of the all the pre-synthesized compounds are shown in Table 1. 3D QSAR Pharmacophore Generation module/Discovery Studio (DS) is employed to build pharmacophore model using hydrogen bond acceptor (HBA), hydrogen bond donor (HBD), hydrophobic (HY) and ring aromatic (RA) and positive ionizable (PI) chemical features^16^. Ten top-scored hypotheses based on the activity values of the training set molecules are produced. The best ten hypotheses are comprised of only two features: HBD and HY. Hypo1 contains two HBD and three HY which establishes the biggest cost difference (69.23), best correlation coefficient (0.91), highest fit value (9.296) and lowest root mean square deviation (RMSD) of 0.88. The fixed and the null cost values are found to be 146.091 and 231.594 respectively where “fixed total cost” is dependent on the sum of the cost components including weight cost, error cost, and configuration cost. Principally, there are two essential contents that are practiced for cost analysis (a) the difference between null and total cost (cost difference) and (b) the difference between the fixed and null costs. The fixed cost corresponds to a cost of the general hypothesis, which ultimately predicts the activity of compounds in the training set with the lowest deviation. On the other hand, the null cost signifies the cost of a hypothesis. The difference between these two costs should be greater than 70 bits to show the over 90% statistical significance of the model. The cost difference must be larger than 60 bits to signify an accurate correlation data. In these results, all the hypotheses have the HBD and HY group, implying that HBD and HY groups play an important role in N N’-Diarylguanidine inhibition. We find that the cost difference between null and fixed cost is 85.50 which is clearly more than 70 bits. All hypotheses have a correlation coefficient of higher than 0.79, but Hypo1 shows the highest correlation coefficient values of 0.91, demonstrating good prediction ability of Hypo1.

**Table 1.**
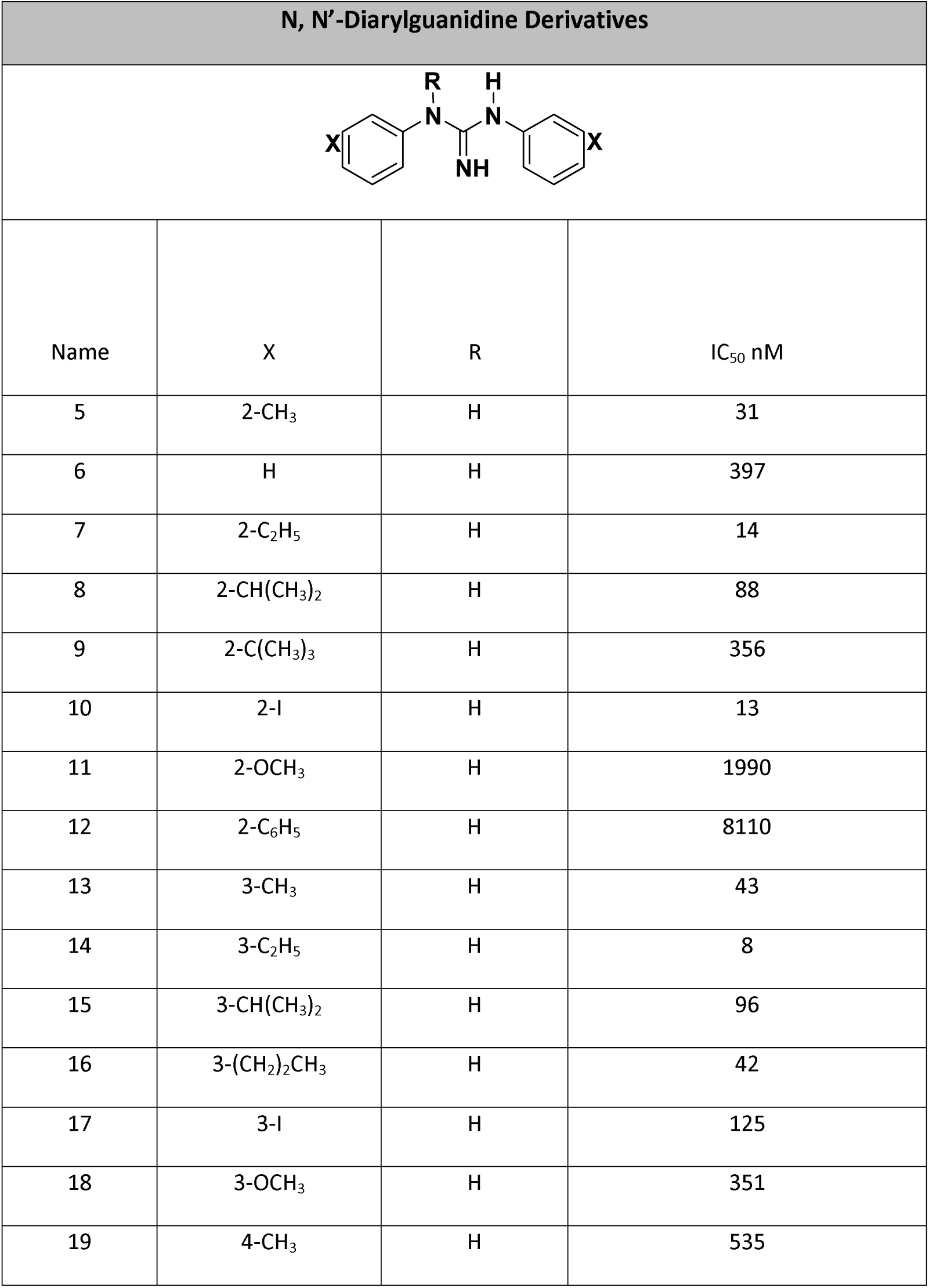
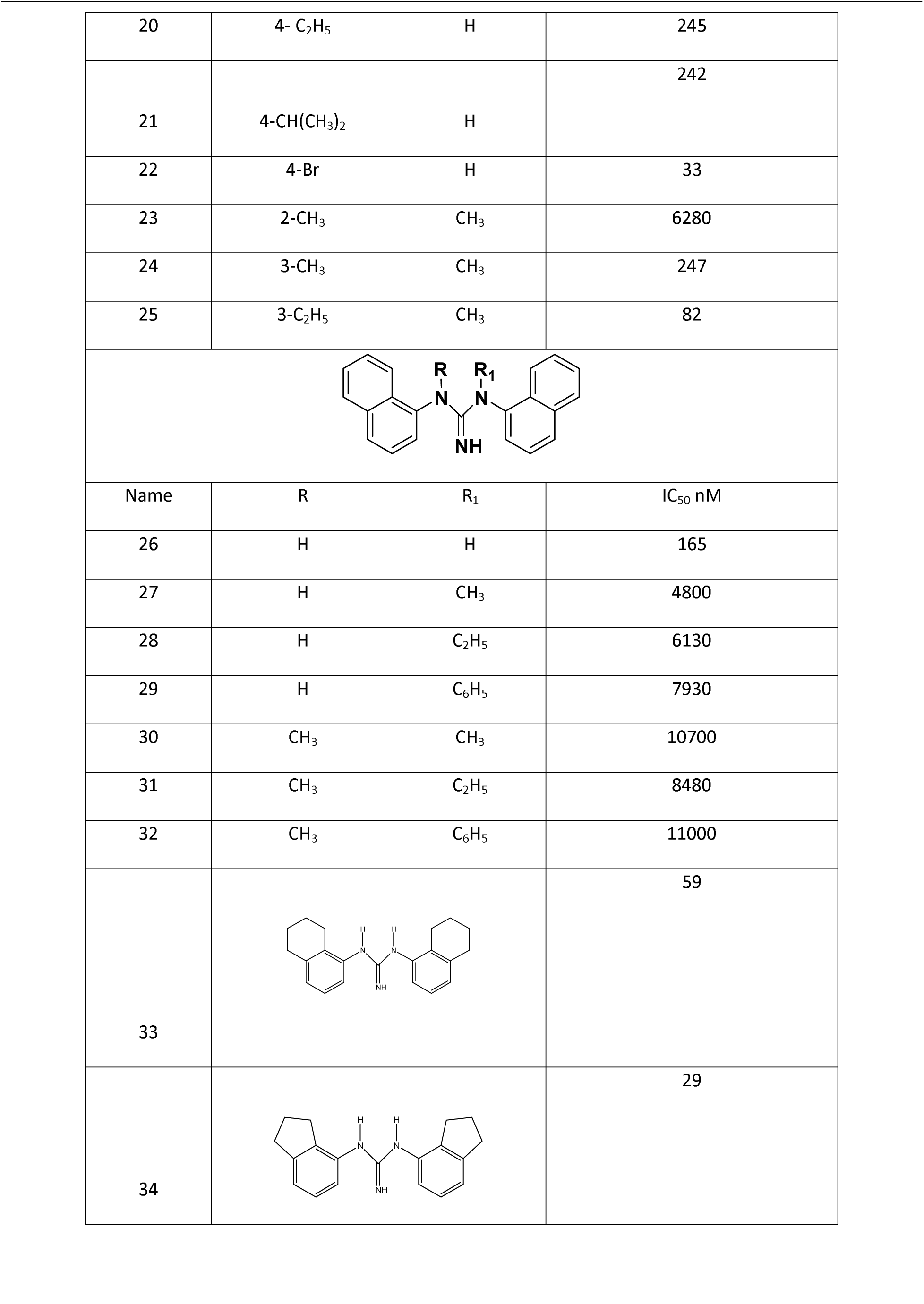
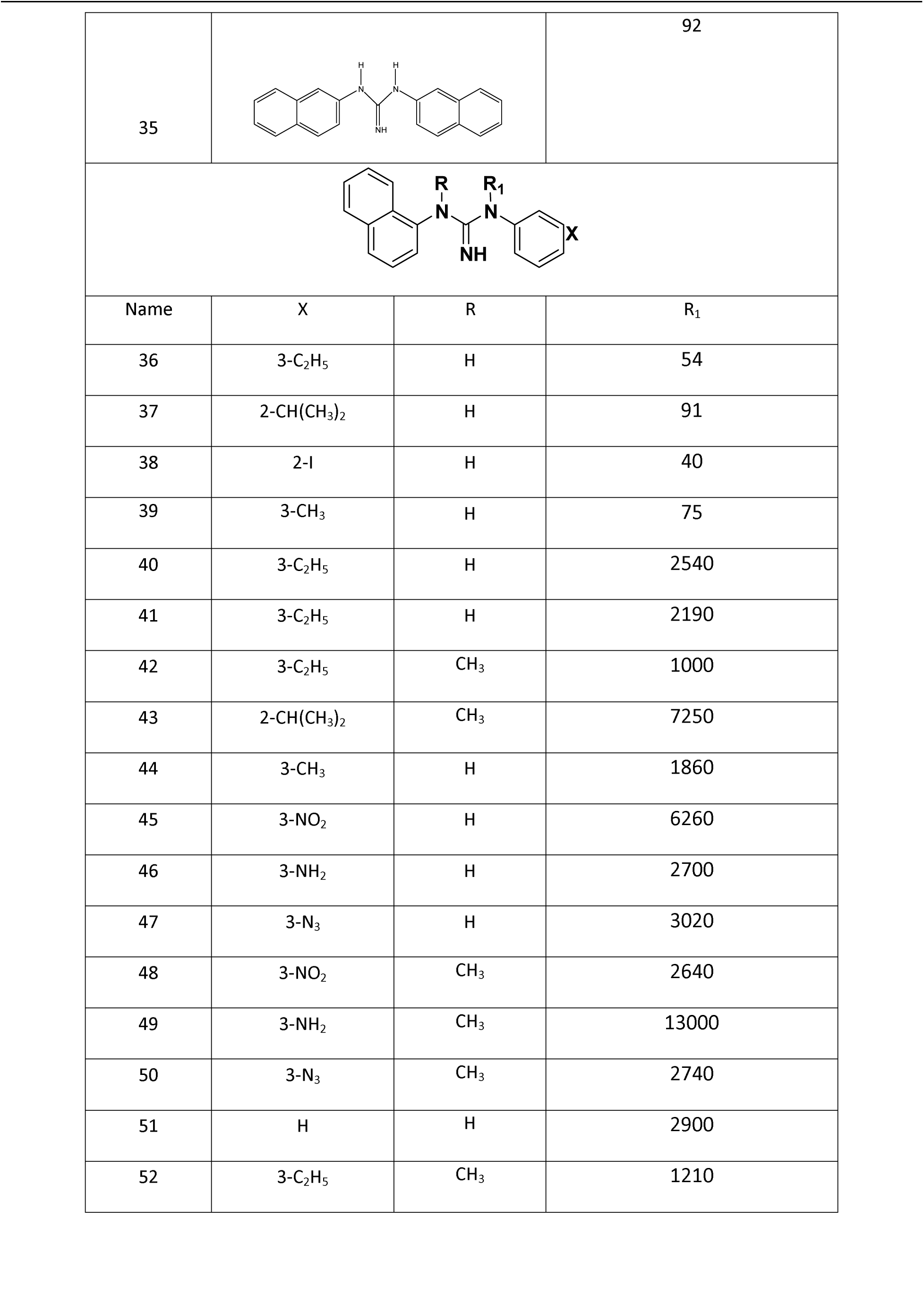
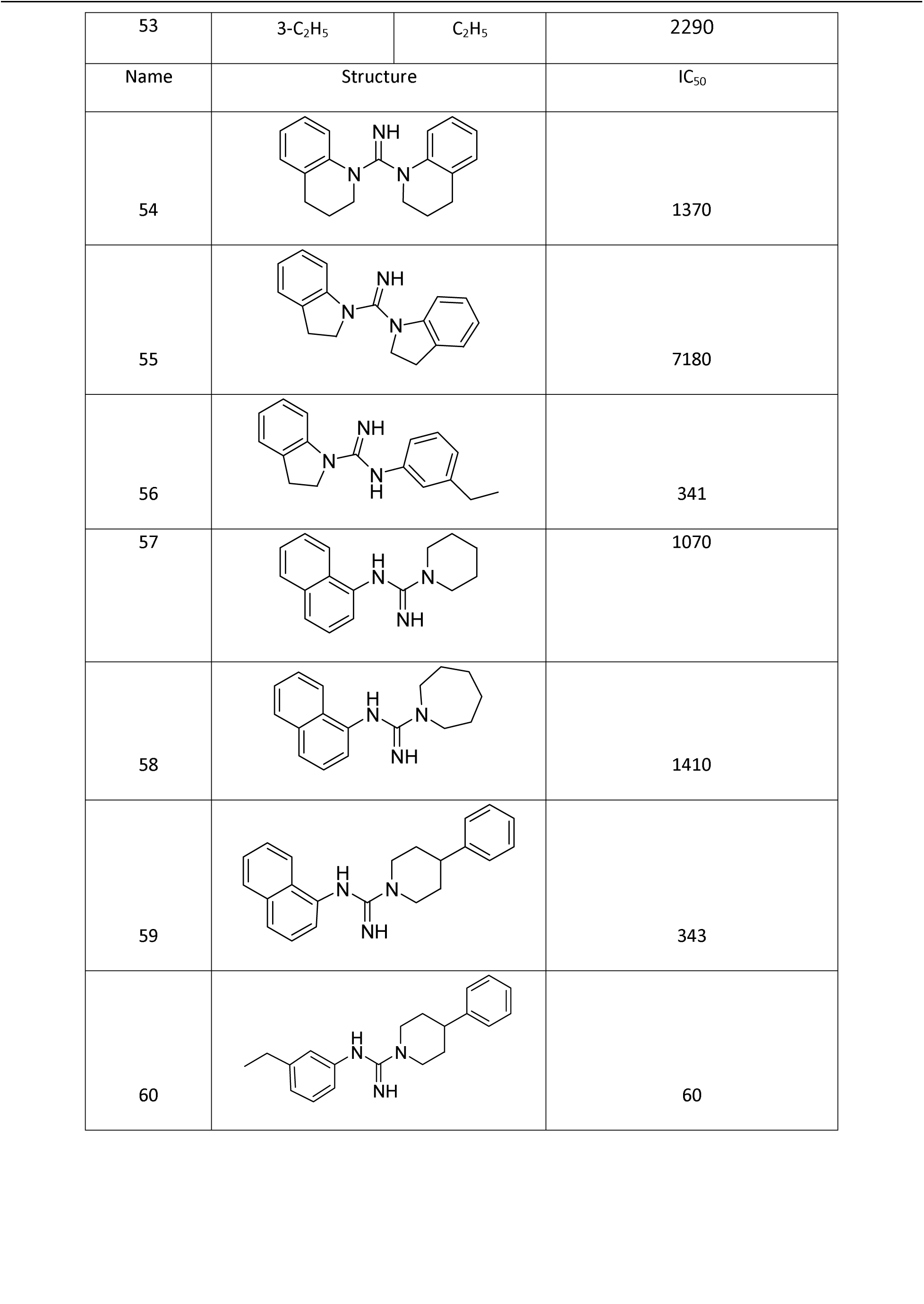
Error! Main Document Only. **Structures and biological activity of N, N’-diarylguanidine derivatives as NMDA receptor antagonists.**

Hypo 1 showed high cost difference and correlation value with lower RMSD values on being evaluated with other hypotheses. Thus, Hypo1 is chosen as the “best hypothesis” and used for other analysis. Fig. 1 shows the chemical features of Hypo1 with its geometric parameters. To verify the prediction accuracy of Hypo1, training set is used and the activity of each compound in training set is estimated by regression analysis. It is observed that “Hypo1” is proficient to assess the activities of compounds. The experimental and estimated activities for training set compounds are shown in Table 2. A plot between the observed activity versus estimated activity demonstrates a good correlation coefficient (r^2^ = 0.83) for training set compounds, indicating the high predictive ability of the pharmacophore as shown in Fig. 2.

**Table 2.**
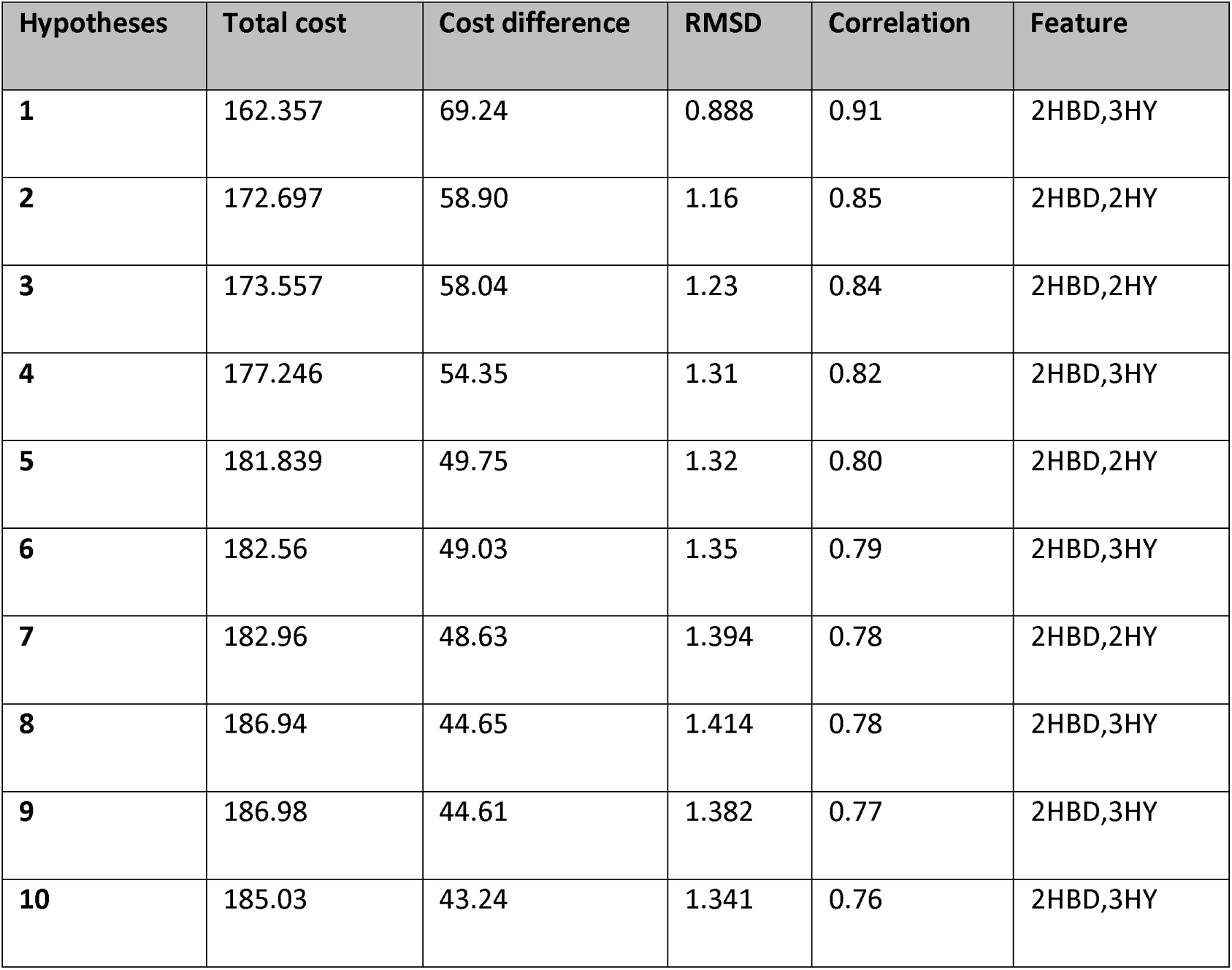
Results of the top 10 pharmacophore hypotheses generated by the hypoGen algorithm.

| Hypotheses | Total cost | Cost difference | RMSD | Correlation | Feature |
| --- | --- | --- | --- | --- | --- |
| 1 | 162.357 | 69.24 | 0.888 | 0.91 | 2HBD,3HY |
| 2 | 172.697 | 58.90 | 1.16 | 0.85 | 2HBD,2HY |
| 3 | 173.557 | 58.04 | 1.23 | 0.84 | 2HBD,2HY |
| 4 | 177.246 | 54.35 | 1.31 | 0.82 | 2HBD,3HY |
| 5 | 181.839 | 49.75 | 1.32 | 0.80 | 2HBD,2HY |
| 6 | 182.56 | 49.03 | 1.35 | 0.79 | 2HBD,3HY |
| 7 | 182.96 | 48.63 | 1.394 | 0.78 | 2HBD,2HY |
| 8 | 186.94 | 44.65 | 1.414 | 0.78 | 2HBD,3HY |
| 9 | 186.98 | 44.61 | 1.382 | 0.77 | 2HBD,3HY |
| 10 | 185.03 | 43.24 | 1.341 | 0.76 | 2HBD,3HY |

**Figure 1.**
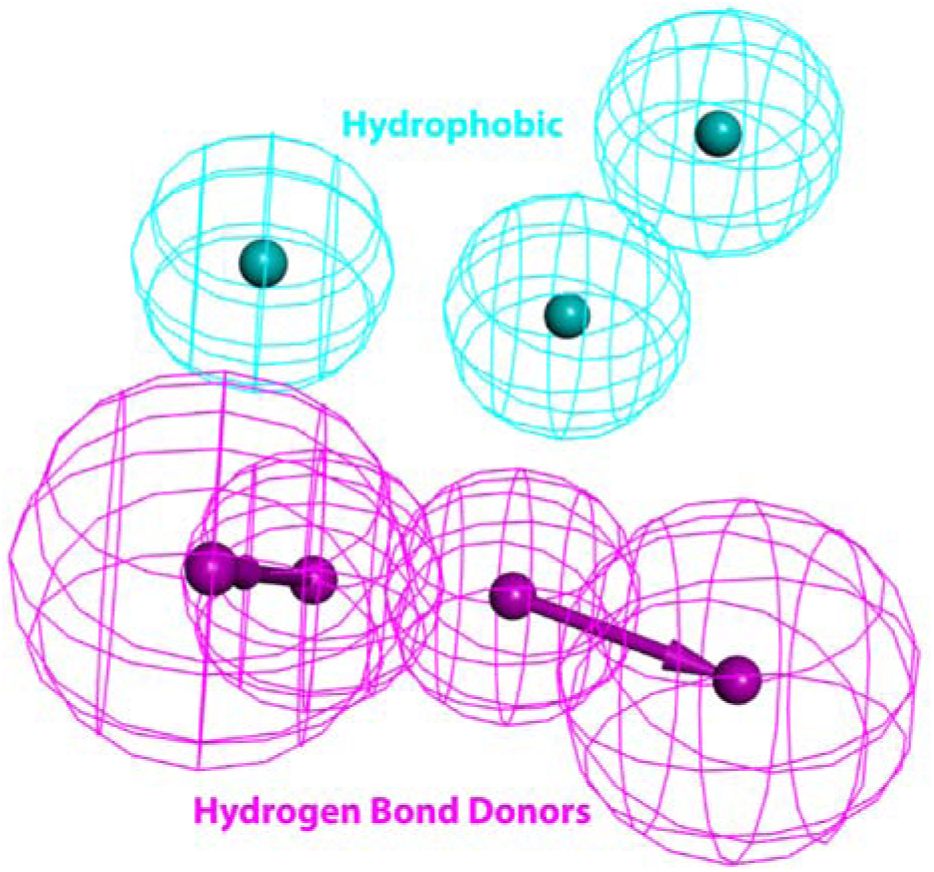
Pharmacophoric features identified from best hypothesis 1.

**Figure 2.**
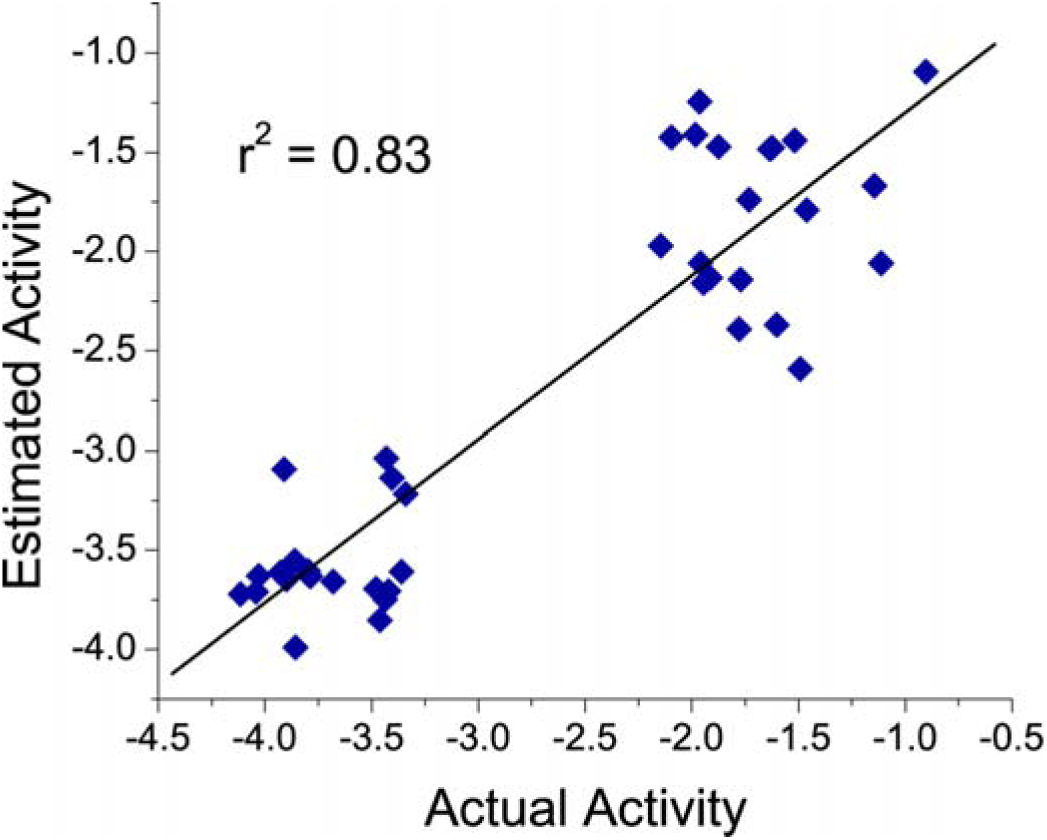
A plot of actual versus estimated biological activity for training set compounds.

### Validation of hypothesis

Validating the generated hypothesis is a vital step in drug development. There are a number of known methods such as (a) preparation of the test set, (b) Fischer’s randomization method and (c) external test set that can assess the quality of pharmacophore. These methods are explained below:

### Fischer randomization method

The foremost reason of this assessment is to confirm the strong correlation amid the chemical structure and biological activities of compounds^38^. The Fischer validation method is applied at a confidence level of 99% to the developed HypoGen model, where 99 random spreadsheets (hypotheses) were generated (Fig. 3). Various pharmacophore hypotheses are generated by randomizing the activity data of the training set compounds with the matching characteristic features and parameters employed to develop the new pharmacophore hypothesis. We observe that none of the resulting hypotheses had a lower cost score than the initial hypothesis which verified that the hypothesis 1 has not been obtained by chance. The data from this validation clearly suggests that all values generated after randomization produces hypotheses with no predictive values similar or near to that of hypotheses1 as shown in Fig. 3. On analyzing the 99 runs, the value of correlation coefficients is found to be relatively low as shown in Fig. 4. The RMS values and the difference between total costs and the null cost are also found to be high which is not considerable for a good model. Hence, this validation procedure provided strong confidence in the chosen pharmacophore model (hypothesis1).

**Figure 3.**
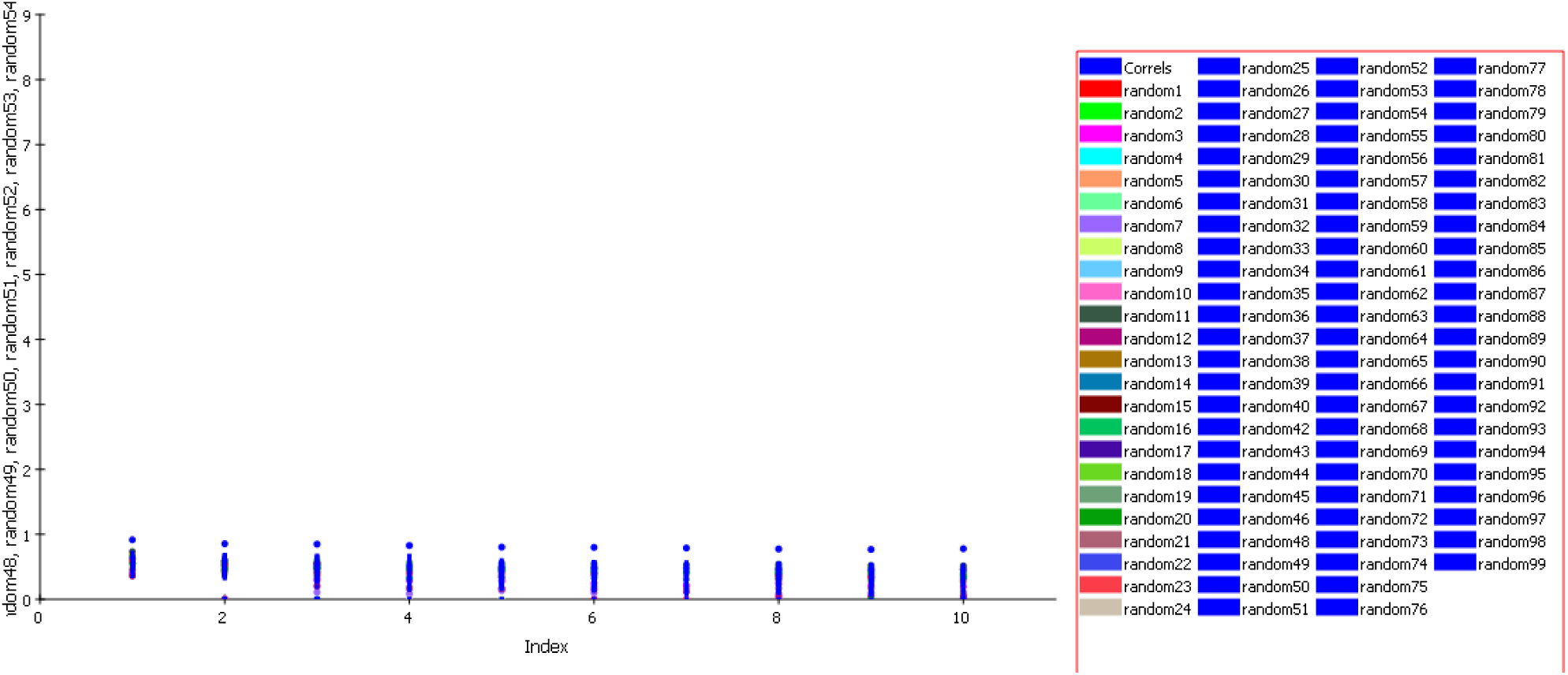
Graph of 99% cat scrambled correlation data. None of the outcome hypotheses has a higher correlation score than the initial (best) hypothesis.

**Figure 4.**
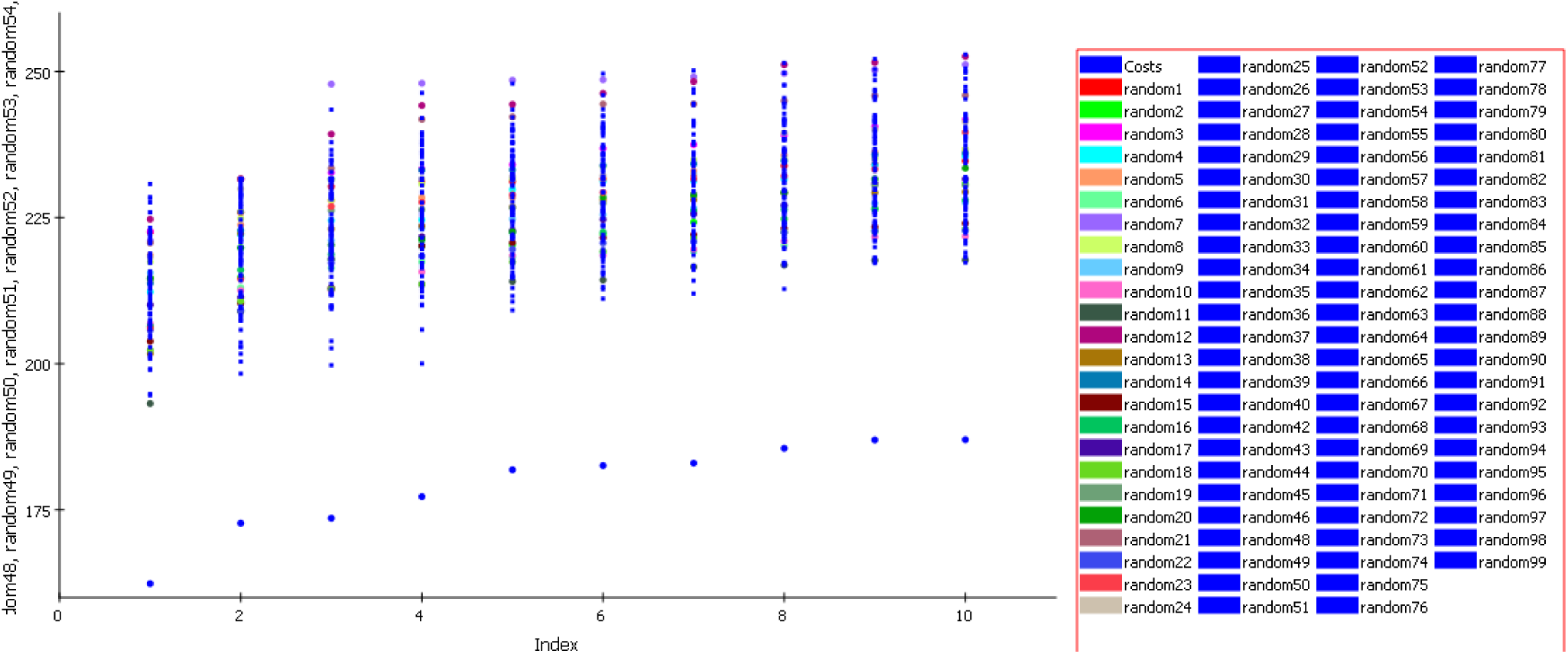
Graph of 99% cat scrambled cost data. None of the outcome hypotheses has a higher correlation score than the initial (best) hypothesis.

### Internal test set validation

In order to confirm whether the generated hypothesis is able to estimate the activity of compounds other than training set compounds, a test set comprising of 19 compounds is used for further validation. The compounds that are not included in model generation are used as the internal test set. All the 19 compounds are mapped onto the generated pharmacophore model using the best fit and conformational energy constraint of 10 kcal mol. Fig. 5 represents a plot between actual and estimated activities of the test set with r^2^ = 0.65. This clearly highlights the good predictive ability of our pharmacophore model.

**Figure 5.**
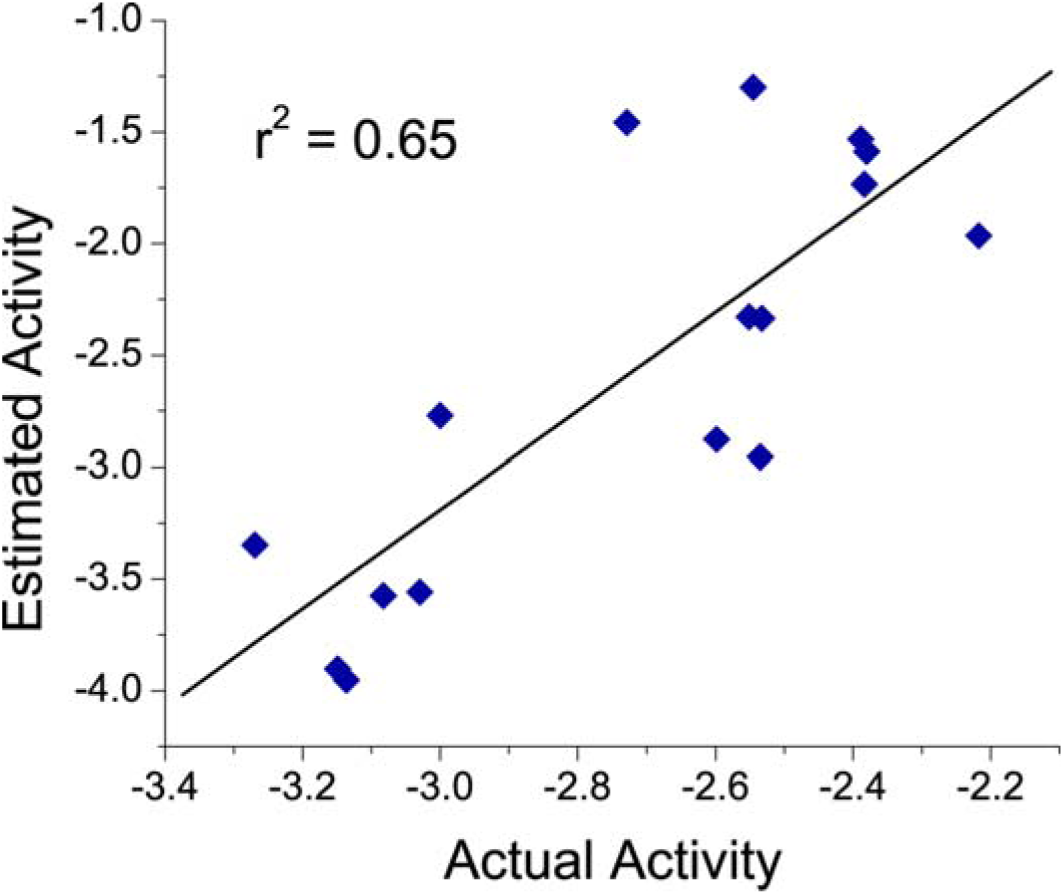
A plot of actual versus estimated biological activity for test set compounds

### External Test Set Validation

The major goal behind the pharmacophore modeling is to utilize them in the hit/lead identification and optimization phases of the drug discovery paradigm. However, predictive quality of the generated model(s) must be evaluated well before its use. External test set validation is one of the best ways to predict the qualities of the generated model. Here, we validate the generated pharmacophore using a structurally diverse external set of NMDA inhibitors. The actual activity of these compounds lies between 0.068 to 2.1 nM. The ten external test set compounds are later mapped onto the generated pharmacophore model. The mapping pattern (fit value) and the difference between the estimated and actual activity of all the compounds is analyzed thoroughly. The mapping of the best fit molecule of the external test set is shown in Fig. 6. The predicted and the actual activity for external test set compounds (**Fig. 7 with r^2^ = 0.87**) testifies the universality of the hypo-1.

**Figure 6.**
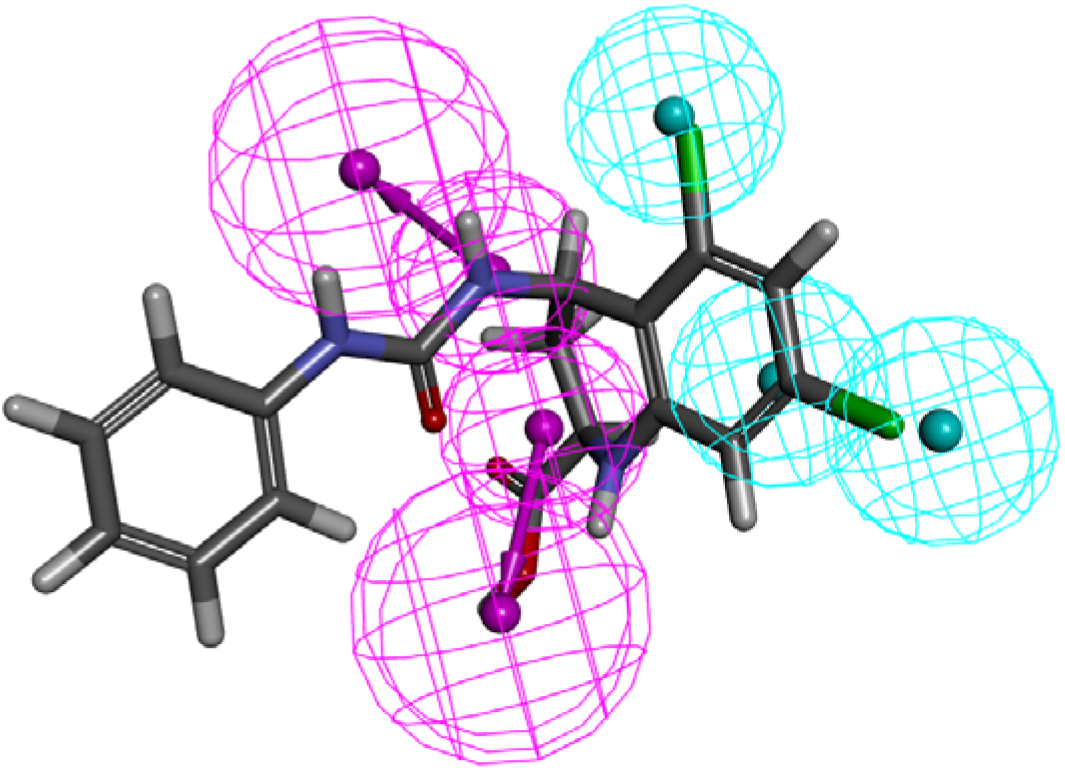
Pharmacophore mapping of most active compound of external test set onto the chosen pharmacophore model (hypothesis 1).

**Figure 7.**
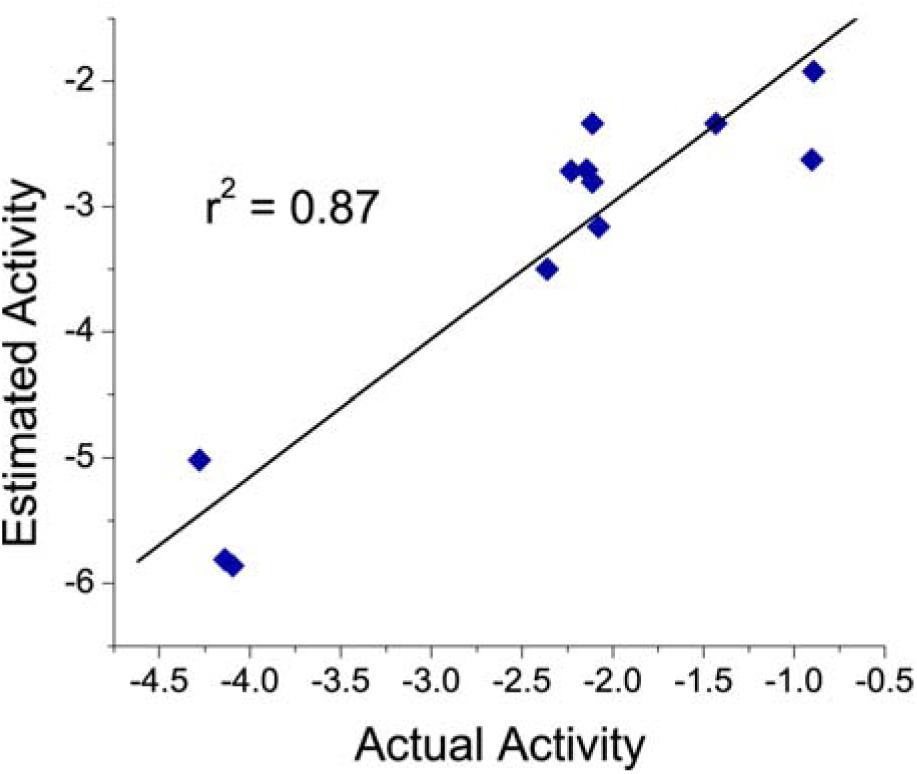
A plot of actual versus estimated biological activity for external test set compounds.

### Database screening

The validated pharmacophore hypothesis, Hypo1, is utilized as a 3D structural query for obtaining compounds from chemical databases including NCI (238 819 compounds) and MayBridge (59 652 compounds). As a result, the first screening resulted in 171 and 299 compounds from NCI and Maybridge respectively^38^. The retrieved hit compounds are filtered on the basis of Lipinski’s violation and the estimated activity values calculated by Hypo 1.

Ten potential compounds are selected after refining a total of 470 hits which display a perfect four feature mapping with a good fit value ranging from 10.18-7.775 respectively and zero Lipinski’s violation.

### Molecular docking studies

Molecular docking is a computational technique that illustrates conformations of compounds in protein binding sites. The main aspects to determine the quality of docking method is docking preciseness, which identifies the correct binding modes of the ligands to the target protein using a docking method. The center of the active site of N-Methyl-D-Aspartate Receptor antagonists is comprised of Tyr214, Thr174, Ser173, Thr116, Arg121, Ser114, Gly172 and His88 which are the significant residues surrounded in this region. 10 (5 Maybridge and 5 NCI) lead compounds retrieved from the databases which satisfy drug like properties as well as one marketed drug (Memantine) are docked in the active site of 3OEK using LibDock software implemented in DS. A total of 300 poses are generated for each compound. Out of those 300, top ten poses are evaluated for each molecule on the basis of LibDock score. We find that HTS 00987 (Maybridge) shows a high LibDock score of 114.714 whereas Memantine (marketed drug) shows the score of 70.16.

The interaction analysis of HTS 00987 reveals that hydrogen present on 2^nd^ and 5^th^ positions of benzodioxol ring is involved in Vander Waals interactions with Tyr214 and Thr174. Amine and hydrogen present on 1-ethylamino-3-methoxy-2-methylpropan-2-ol shows hydrogen bond and Vander Waals interactions with Pro170, His88 and Lys87 respectively as shown in Fig.8 (A). Interaction analysis of reference drug memantine reveals that amine group present on adamantan-1-amine ring is involved in hydrogen bond interactions with Ser131, Tyr282, Gly264 and Ser260. Methyl present on the 3^rd^ and 5^th^ position of amine ring showed Vander Waals interactions with Asp283, His127 and Arg292 as shown in **Fig.8 (B)**. Another Carbon present at the second and third positions of methoxyphenyl amino group is interacting with Tyr214 with an interfeature distance of 1.031. The interaction analysis of memantine shows hydrogen bond interactions with amino acid residues His88, Tyr214 and Gly13 Fig.8 (B).

**Figure 8.**
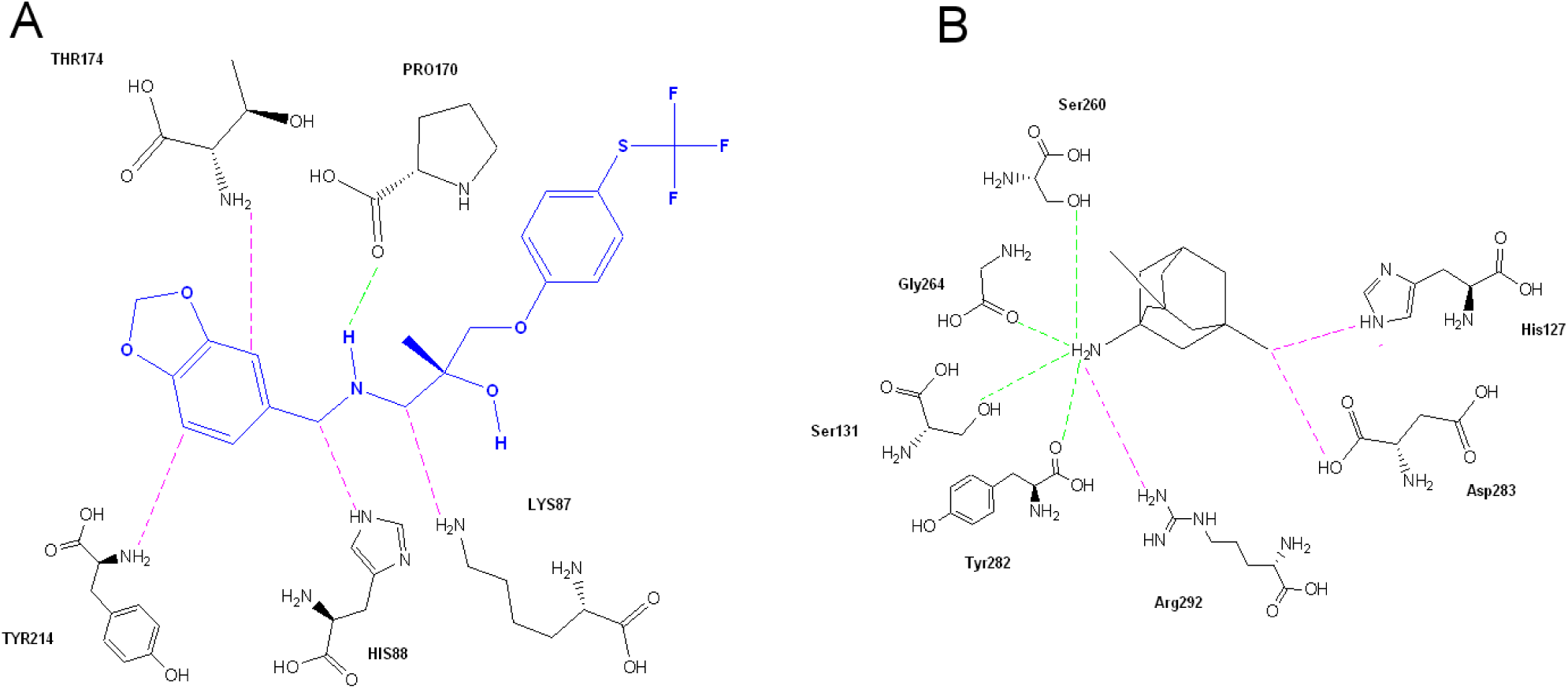
(A) Interaction of HTS 00987 with Tyr214, Thr174, Pro170, His88 and Lys87 in the active site. [Green dotted lines represent Hydrogen bond interactions; Pink dotted lines represent Vander Waals interaction] (B) Interaction of Memantine with Ser131, Tyr282, Gly264, Ser260, Asp283, His127 and Arg292 in the the active site. [Green dotted lines representing Hydrogen bond interactions; Pink dotted lines representing Vander Waals interaction]

In view of good fit values, estimated activity, drug-likeness, docking score and availability for procuring, HTS 00987 is checked for novelty by employing pairwise tanimoto similarity indices using “Find Similar Molecules by Fingerprint” protocol in Discovery Studio. HTS 00987 shows low tanimoto similarity indices of 0.014 to all the structures of known NMDA inhibitors confirming their novelty^39^. Hence, in the course of this research attempt, we discover one druggable potent N-Methyl-D-Aspartate Receptor antagonist which can be further explored in clinical trials.

### Analysis of the *in-vivo* studies

After rigorous validation, HTS 00987 is subjected to *in-vivo* studies. The experimental design is planned such that the effect of all the standard and test drugs can be evaluated after 7 and 14 days against diazepam induced amnesia. The cognitive parameters are evaluated 30 minutes after the administration of inducing agent (Diazepam) using an eight arm radial maze. On day 7 and 14, the lead compound (HTS 00987) demonstrates a significant increase in number of entries and duration in baited arms (**Day 7**: M = 17.25, S.D = 1.5; **Day 14**: M = 21.5, S.D = 2.3, p < 0.05) as compared to memantine (**Day 7**: M = 14.75, S.D = 2.38; **Day 14**: M = 16.85, S.D = 2.21) as shown in Fig.9 (A). Similarly, we observe an increase in duration in baited arms (**Day 7**: M = 364.37, S.D = 5; **Day 14**: M = 386.72, S.D = 4.1, p < 0.05) when compared to memantine (**Day 7:** M = 234.45, S.D = 4.01; **Day 14**: M = 369.19, S.D = 5, p < 0.05) Fig.9 (B). The results also show a significant decrease in RME (**Day 7**: M = 15.75, S.D = 3.61; **Day** 14: M = 7.25, S.D = 1.88, p < 0.05), and WME (**Day 7**: M = 4, S.D = 0.62; **Day 14**: M = 3.5, S.D = 0.86, p < 0.05) when compared with memantine [**RME (Day 7**: M = 16, S.D = 5.23; **Day 14**: M = 13.25, S.D = 4.12), **WME** (**Day 7**: M = 6, S.D = 1.12; **Day 14**: M = 5.25, S.D = 2.93)] which is clearly seen in Fig.9 (C). Another major cognitive parameter, percent choice is also calculated and it is observed that HTS 00987 demonstrates a high percent choice (**Day 7**: M= 54.14, S.D= 1.05; **Day 14**: M= 55.98, S.D= 2.01, p<0.05) when compared with memantine (**Day 7**: M= 47.5, S.D= 1.11; **Day 14**: M= 52.5, S.D= 1.89, p<0.05) as shown in Fig.9 (D).

**Figure 9.**
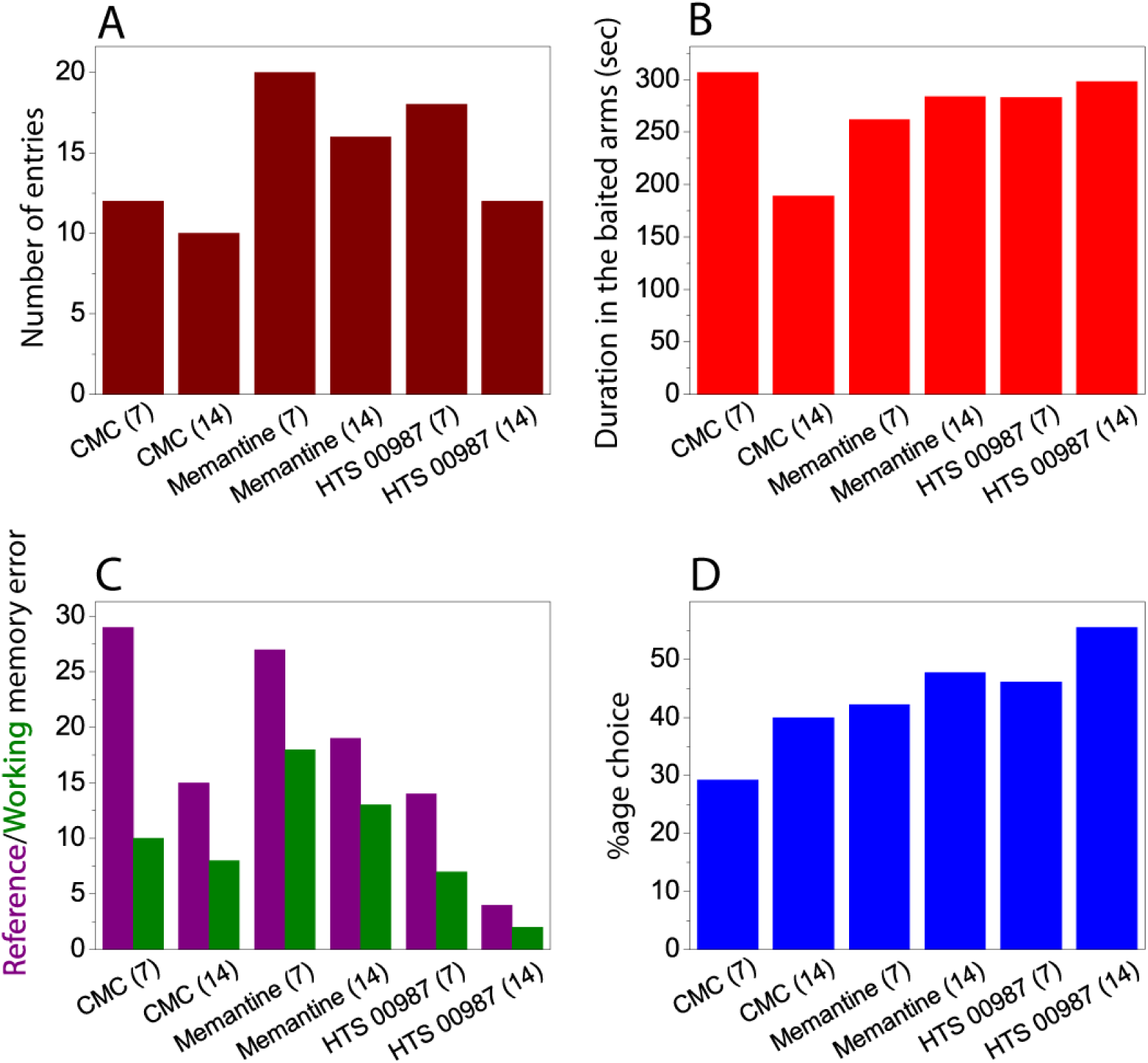
The effects of CMC, memantine and HTS 00987 on an eight-arm radial maze in diazepam-induced amnesia in mice: **(A)** Number of entries in baited arm. **(B)** Duration in baited arms *(in sec)* **(C)** Reference/Working Memory Errors **(D)** Percentage of correct choices.

In summary, HTS 00987 shows a remarkable increase in the number of entries in the baited arm, duration in baited arms, % correct choice and a significant decrease in RME and WME in diazepam-induced amnesia in mice as compared to memantine during week 1 and surprisingly its repeated administration also showed a noteworthy increase in memory. In view of the aforementioned results, it can be concluded that HTS 00987 has shown good neuroprotection properties and the lead compound can be subjected to clinical trials for their development as novel drugs.

## Conclusion

The virtual screening protocol using the highly validated pharmacophore model leads to the identification of several druggable compounds, one of which is experimentally validated. In the presented results, the lead compound has shown a good *in vivo* potency. The compound identified has a potential to assist the development of novel neuroprotective agents for the treatment of Alzheimer’s disease. Moreover, the identified compounds can serve as a template for designing new NMDA receptor antagonists. In conclusion, our study clearly shows that if elucidated and validated properly, the ligand-based pharmacophore can be a powerful source for identification of novel leads from chemical compound databases.

## Ethics approval and consent to participate

All experiments performed were approved by the Institutional animal ethical committee.

## Author Contributions

MS performed the whole experiments. SP has made major contribution to conception and design of the experiments. SS contributed in the pharmacological studies. AM, AS and AKJ supported in analyzing the results. MS wrote the manuscript.

## References

1 Francesco G S, Boyd, L H, Bruce M B, Philip L. N, Kenneth T S, John H K H, Steven W, Ian A (1992) 3-(2-Carboxyindol-3-yl) propionic Acid-Based Antagonists of the N-Methyl-D-aspartic Acid Receptor Associated Glycine Binding Site. J Med Chem 35: 1791–1799

2 Cull-Candy S, Brickley S, Farrant M (2001) NMDA receptor subunits diversity, development and disease. Curr Opin Neurobiol 11: 327–335

3 Kew J N, Trube G, Kemp J A (1998) State-dependent NMDA receptor antagonism by Ro8-4304, a novel NR2B selective, non-competitive, voltage-independent antagonist. Br J Pharmacol 3: 463 –472

4 Chenard B L, Menniti F S (1999) Antagonists selective for NMDA receptors containing the NR2B subunit. Curr Pharm Des 5: 381–404

5 Bobich J A, Zheng Q, Campbell A (2004) Incubation of nerve endings with a physiological concentration of αβ 1–42 activates CaV2.2 (N-Type)-voltage operated calcium channels and acutely increases glutamate and noradrenaline release. J Alzheimers Dis 6: 243–255

6 John F K, Robert N B, Michael W, James B F, Philip N H, Stephen M S, Alfred C S, Steven F, Charles F S, Craig J, Eckard W (1989) Synthesis and characterization of a series of diarylguanidines that are noncompetitive N-methyl-D-aspartate receptor antagonists with neuroprotective properties. Proc Natl Acad Sci U S A 86: 5631–5635

7 Sharma M, Mittal A, Singh A, Jainarayanan A, Paliwal S (2018) Identification of novel and structurally diverse N-Methyl-D-Aspartate Receptor Antagonists: Successful Application of Pharmacophore Modeling, Virtual Screening and Molecular Docking. bioRxiv doi:https://doi.org/10.1101/314914

8 Gunner O A (2000) Pharmacophore, perception, development, and use in drug design. Uni Intl L: S Diego 17–20

9 Li H, Sutter J, Hoffman R (1999) HypoGen: An automated system for generating 3D predictive pharmacophore models. In: Pharmacophore Perception, Development, and Use in Drug Design. Intl Uni L: La Jolla, CA: 171–189

10 Kurogi Y, Guner O. F (2001) Pharmacophore modeling and three dimensional database searching for drug design using catalyst. Curr. Med. Chem 22:1035–1055

11 Meganathan C, Sakkiah S, Lee Y, Narayanan J V, Lee K.W (2013) Discovery of potent inhibitors for interleukin-2-inducible T-cell kinase: structure-based virtual screening and molecular dynamics simulation approaches. J Mol Model. 19: 715–726

12 Li H, Sutter J, Hoffmann R D (2000) Pharmacophore perception, development, and use in drug design, IUL Biotechnology Series. Güner OF, editor. La Jolla, CA: International University Line;. HypoGen: An automated system for generating 3D predictive pharmacophore models 171–189.

13 Kurogi Y, Güner OF (2001) Pharmacophore modeling and three-dimensional database searching for drug design using catalyst. Curr Med Chem 8:1035–1055.

14 A.K. Debnath (2002) Pharmacophore Mapping of a Series of 2,4-Diamino-5-deazapteridine Inhibitors of Mycobacterium avium Complex Dihydrofolate Reductase. J. Med. Chem. 45: 41–53

15 Pal M, Paliwal S (2012) In silico identification of novel lead compounds with AT1 receptor antagonist activity: successful application of chemical database screening protocol. Org Med Chem Lett 2: 1–7

16 Sakkiah S, Thangapandian S, John S, Kwon Y J, Lee K W (2010) 3D QSAR pharmacophore based virtual screening and molecular docking for identification of potential HSP90 inhibitors. Eur J Med Chem 45: 2132–2140

17 Mittal A, Paliwal S, Sharma M, Singh A, Sharma S, Yadav D (2014) Pharmacophore based virtual screening, molecular docking and biological evaluation to identify novel PDE5 inhibitors with vasodilatory activity. Bioorg Med Chem Lett 24:3137–41

18 Acharya B N, Kaushik M P (2007) Pharmacophore-based predictive model generation for potent antimalarials targeting haem detoxification pathway. Med Chem Res 16:213–229

19 Bhattacharjee A, Mylliemngap B J, Velmurugan D (2012) 3D-QSAR studies on fluroquinolones derivatives as inhibitors for tuberculosis. Bioinformation 8:381–387

20 Mehta H, Khokra S L, Arora K, Kaushik P (2012) Pharmacophore mapping and 3D-QSAR analysis of Staphylococcus Aureus Sortase a inhibitors. Der Pharma Chemica 4:1776–1784

21 Chandra K U, Haripriya M, Shaik M (2010) 2D QSAR, pharmacophore and docking studies of mycobacterium tuberculosisenoyl acyl carrier protein reductase inhibitors. J Glo Pharm Tech 2:73–89

22 Walters W P, Stahl M T, Murcko M A (1998) Virtual screening –an overview. Drug Discov Today 3:160–178.

23 Hou T, Xu X (2004) Recent development and application of virtual screening in drug discovery: An overview. Curr Pharm Des 10:1011–1033

24 Anderson A C, Wright D L (2005) The design and docking of virtual compound libraries to structures of drug targets. Curr Comput Aided Drug Des 1:103–127

25 Daisy P, Suveena S, Lilly V (2011) Molecular docking of medicinal compound Lupeol withautolysin and potential drug target of UTI. J Chem Pharm Res 3:557–562

26 Sharma M, Jainarayanan (2018) AchE-OGT dual inhibitors: Potential Partners in Handling Alzheimer’s Disease. Biorxiv: 10.1101/303040

27 Fengbo Wu, Ting Xu, Gu He, Ouyang L, Han B, Peng C,Song X, Xiang M (2012) Discovery of novel focal adhesion kinase inhibitors using a hybrid protocol of virtual screening approach based on multicomplex-based pharmacophore and molecular docking. Int J Mol Sci 13:15668–15678

28 Meng X Y, Hong X Z, Cui M M M (2011) Molecular Docking: A powerful approach for structure-based drug discovery. Curr Comput Aided Drug Des 146–157

29 Liu P, Bilkey D K (1999) The effect of excitotoxic lesions centered on the perirhinal cortex in two versions of the radial arm maze task. Behav Neural Biol 113: 672–682

30 Suzuki S W, Augerinos G, Black B A (1980) Stimulus control of spatial behavior on the eight-arm maze in rats. Learning and Motivation 11:1–18

31 Kirti S K, Kasture S B, Mengi S A (2010) Efficacy study of Prunus Amygdalus(almond) nuts in scopolamine-induced amnesia in rats. Indian J Pharmacol 42:168–173

32 Sudesh P, Manish K S, Promila P, Akshay A (2008) Bacopa monniera exerts antiamnesic effect on diazepam-induced anterograde amnesia in mice. Psychopharmacology 200: 27–37

33 Olthof A, Sutton J E, Slumskie S V D,’Addetta J, Roberts W (1999) A. In search of the cognitive map: Can rats learn an abstract pattern of rewarded arms on the radial maze? J Exp Psychol Anim Behav Process 25:352–362

34 O’Keefe J, Nadel L (1978) The hippocampus as a cognitive map. Oxford: Oxford University Press

35 Brown, M F, Wintersteen J (2004) Spatial patterns and memory for locations. Learning & Behavior 32:391–400

36 Kay C H, Hunt M (2010) Differential effects of MDMA and scopolamine on working versus reference memory in the radial arm maze task. Neurobiol Learn Mem 2:151–156

37 Castner S A, Goldman R, Patricia S W, Graham V (2004) Animal models of working memory: Insights for targeting cognitive dysfunction in schizophrenia working memory: The core cognitive deficit in schizophrenia. Psychopharmacology 1:111–125

38 John S, Thangapandian S, Sakkiah S, Lee K W (2011) Potent bace-1 inhibitor design using pharmacophore modeling, in silico screening and molecular docking studies. BMC Bioinformatics 12(Suppl 1):S28.

39 Willett P, Barnard J M, Downs G M (1998) Chemical similarity searching. J Chem Inf Comput Sci 38:983–996.

